# Exploring Extreme Signaling Failures in Intracellular Molecular Networks

**DOI:** 10.1101/2021.04.20.440674

**Authors:** Mustafa Ozen, Effat S. Emamian, Ali Abdi

## Abstract

Developing novel methods for the analysis of intracellular signaling networks is essential for understanding interconnected biological processes that underlie complex human disorders. A fundamental goal of this research is to quantify the vulnerability of a signaling network to the dysfunction of one or multiple molecules, when the dysfunction is defined as an incorrect response to the input signals. In this study, we propose an efficient algorithm to identify the extreme signaling failures that can induce the most detrimental impact on the physiological function of a molecular network. The algorithm basically finds the molecules, or groups of molecules, with the maximum vulnerability, i.e., the highest probability of causing the network failure, when they are dysfunctional. We propose another algorithm that efficiently accounts for signaling feedbacks in this analysis. The algorithms are tested on two experimentally verified ERBB and T cell signaling networks. Surprisingly, results reveal that as the number of concurrently dysfunctional molecules increases, the maximum vulnerability values quickly reach to a plateau following an initial increase. This suggests the specificity of vulnerable molecule (s) involved, as a specific number of faulty molecules cause the most detrimental damage to the function of the network. Increasing a random number of simultaneously faulty molecules does not further deteriorate the function of the network. Such a group of specific molecules whose dysfunction causes the extreme signaling failures can better elucidate the molecular mechanisms underlying the pathogenesis of complex trait disorders, and can offer new insights for the development of novel therapeutics.

## INTRODUCTION

Cellular functions are largely regulated by signaling events within the complex intracellular molecular networks [1-3]. Signals are typically transmitted from the cell membrane to the nucleus via intracellular signaling networks, to regulate some target molecules and alter different cellular functions. Intracellular signaling networks have been studied to address a variety of different questions [1-8].

An important application of the studies on intracellular signaling networks is in target discovery and drug development. In the area of targeted therapies in pharmaceutical industry, a wide variety of highly effective therapeutics have been successfully developed that can target the function of a selective set of molecules within the complex intracellular signaling networks. Fault diagnosis is a platform technology for finding such selective targets by using computational and systems biology techniques that have been developed and optimized over the past few years [9-12]. The main purpose of such technology developments is to understand how vulnerable the entire network is to the dysfunction of each molecule, or a specific group of molecules. In the realm of fault diagnosis technology development, the dysfunctional state of a molecule can be defined as the failure to respond correctly to the input signals, which may further propagate into incorrect responses at the output of the network. For this analysis, we have defined the vulnerability level of a molecule as the probability of having incorrect network responses when that specific molecule is dysfunctional. Vulnerability analysis can be performed for the dysfunction of a single molecule, as well as a group of molecules. The importance of the latter can be attributed to the widely known observations that the most common human disorders, such as cancer or schizophrenia, are reported to be associated with the dysfunction of multiple molecules rather than a specific single molecule [13,14]. This is in contrast to some rare genetic disorder when a single molecule or a genetic mutation is sufficient to cause the pathology [15]. By computing the vulnerability level of a molecule or a group of molecules, one can identify and rank the key molecular components for gaining a desired response, and then compute how much they contribute to the failure of the network. From the drug development stand points, the highly vulnerable molecules can be used as the molecular targets of novel therapeutics.

To analyze a molecular network, first a biologically relevant model needs to be adopted. Here we consider the class of discrete models such as Boolean models [1-6,8,16]. They are particularly useful as they do not need detailed kinetic information and still provide certain biologically relevant insights and predictions. Compared to continuous differential equations models [7], discrete models do not require the knowledge of many mechanistic details and numerous kinetic parameters, and are more appropriate for our study in this paper.

The main goal of this paper is to develop a systematic method to explore the extreme failures of intracellular signaling networks. We define the extreme or the worst possible signaling failure as a pathological phenomenon that results in the highest probability of network failure, where the network failure is defined as the level of departure of the network response from its normal or expected response. The said pathological phenomenon is characterized to be emerged from the presence of one or more dysfunctional molecules in the network. It is conceivable that different individual dysfunctional molecules can result in different levels of probabilities of the network failure. It is not clear, however, what happens if two or more molecules are concurrently dysfunctional, and if the network failure probability increases with the number of simultaneously dysfunctional molecules or not. It is also of interest to have an efficient algorithm to determine the maximum possible network failure probability, over the large number of all possible groups of dysfunctional molecules. The computational complexity of an exhaustive search approach for a network with *K* molecules is extremely high, in the order of *K*^*K*/2^, which is unmanageable as *K* increases. Here we introduce a computationally efficient algorithm with a much less running time in the order of *K*^3^, for identifying the worst possible signaling failures, considering multiple dysfunctional molecules. This study is particularly important in the context of complex disorders with unknown molecular sources, where more than one molecule is observed to be involved in the pathology [13,14].

In this study we first present our extreme signaling failure analysis results on the ERBB signaling network of [17] and the T cell signaling network of [2]. Then, we analyze the computational complexity of the proposed algorithm and compare it with the exhaustive search. Afterwards, we provide the details of the vulnerability analysis equations and the proposed extreme signaling failure analysis algorithm in Methods, Sections A and B, respectively. Moreover, we propose another algorithm in Methods, Section C, that determines the number of time points (clock cycles) needed for network analysis and simulation while computing the vulnerability levels, so that we prevent running network simulations longer than what is needed. We present the results of this algorithm, when applied to the ERBB and the T cell signaling networks, in Methods, Sections D and E, respectively, and finally conclude the paper with some concluding remarks.

## RESULTS OF THE EXTREME SIGNALING FAILURE ANALYSIS

The extreme signaling failure algorithm was applied to two networks: a small signaling network that regulates the transmembrane tyrosine kinase ERBB [17], a therapeutic target in breast cancer, and a larger T cell signaling network [2] involved in a variety of immune system response. Results reveal the impact of the number of dysfunctional molecules in causing the extreme signaling failure.

### A. Extreme Signaling Failure Study of ERBB Network

The ERBB network (Figure 1) has one input and one output. The input molecule is the Epidermal Growth Factor (EGF), whereas the output molecule is the Retinoblastoma Protein (pRB). This network has been studied in the context of breast cancer and understanding the mechanism of action of few drug molecules [17]. Equations that specify how the activity of each molecule is regulated by its inputs are listed in Table 1. To model a dysfunctional molecule, we assume its activity state is either stuck-at-0, SA0, or stuck-at-1, SA1, each with a probability of 1/2 [9].

**Figure 1.**
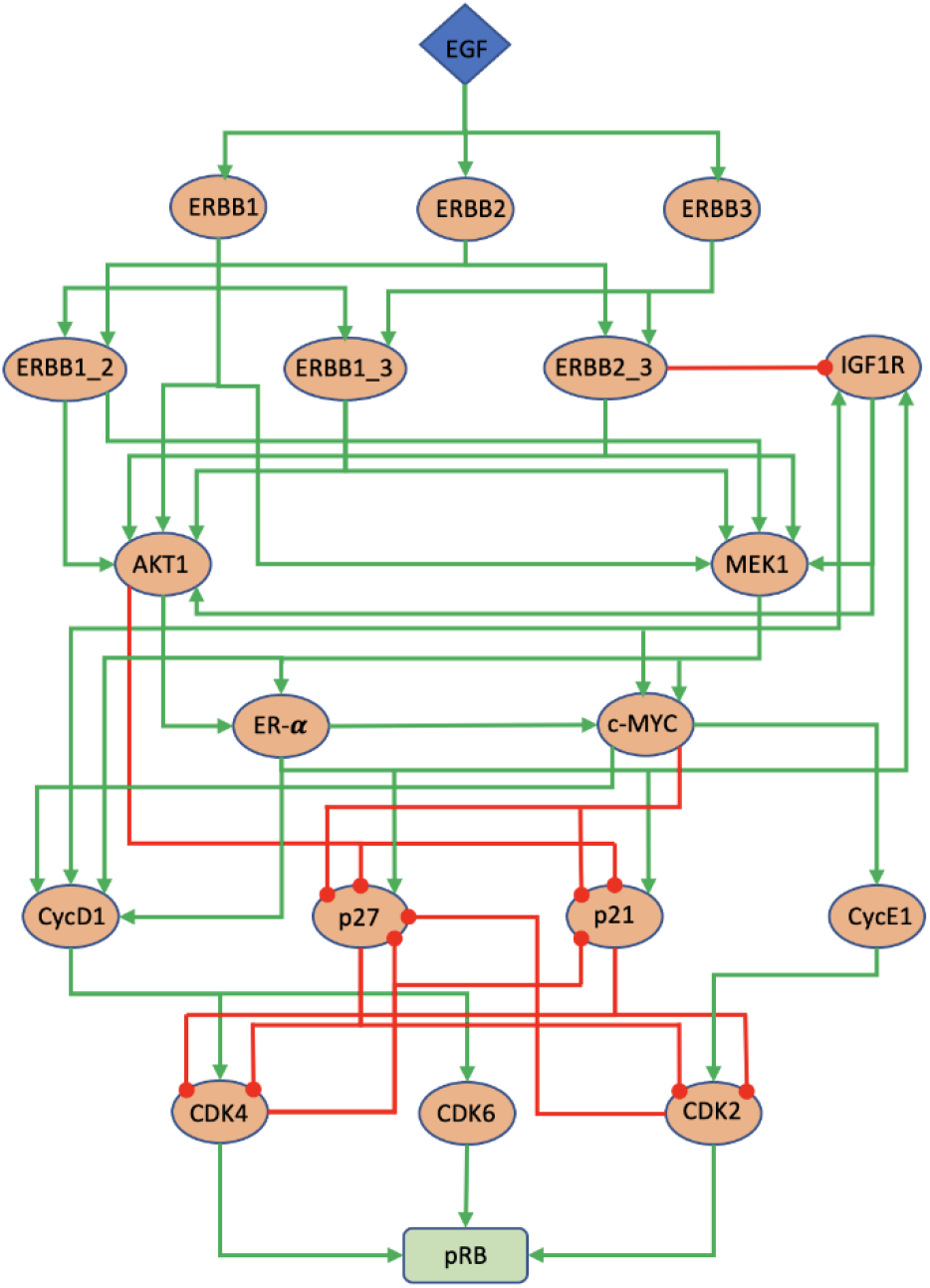
The experimentally-verified ERBB signaling network reproduced from the study of Sahin *et al* [17]. The green arrows represent activatory interactions and the red circle-ended edges represent inhibitory interactions. The input and output nodes represent EGF and pRB, respectively.

**Table 1.**
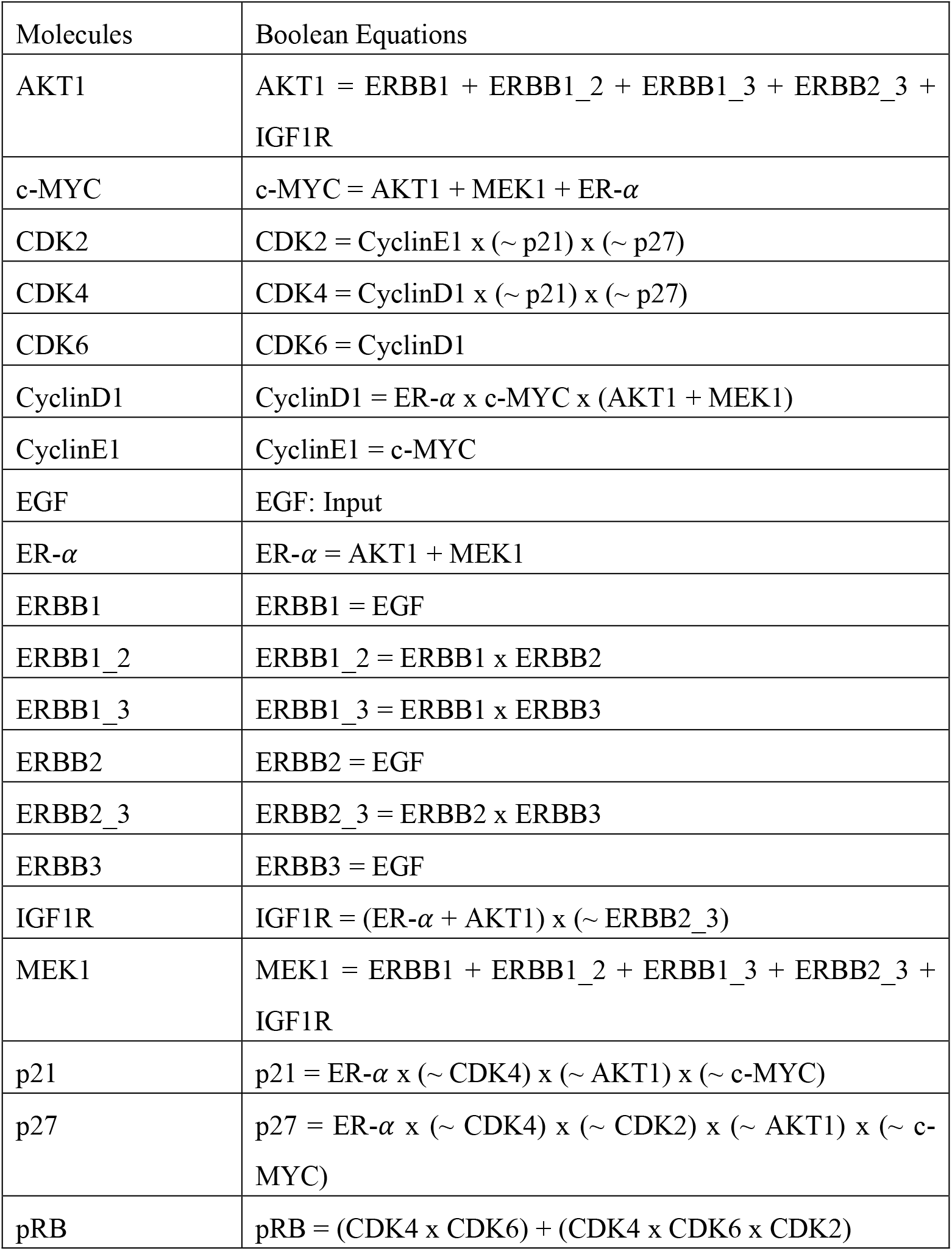
The Boolean equations for the ERBB signaling network (Figure 1) provided in [17]. In the equations, “x” is used for the AND operation, “+” is used for the OR operation, and “∼” is used for the NOT operation.

Let *N* be the number of molecules that are simultaneously faulty, i.e., dysfunctional, in the network. According to the developed vulnerability analysis equations (Methods, Section A) and using the proposed extreme signaling failure algorithm (Methods, Section B), the network maximum vulnerability is computed for each *N* (Figure 2). Here *N* varies from 1 to 18, since the number of intermediate molecules in the network (Figure 1) is 18. For any *N*, the network maximum vulnerability is the highest probability of network failure, where the network failure is defined as the level of departure of the network response from its normal values. In other words, the network maximum vulnerability for a given *N* is a parameter that quantifies the extreme possible signaling failure when there are *N* faulty molecules in the network.

**Figure 2.**
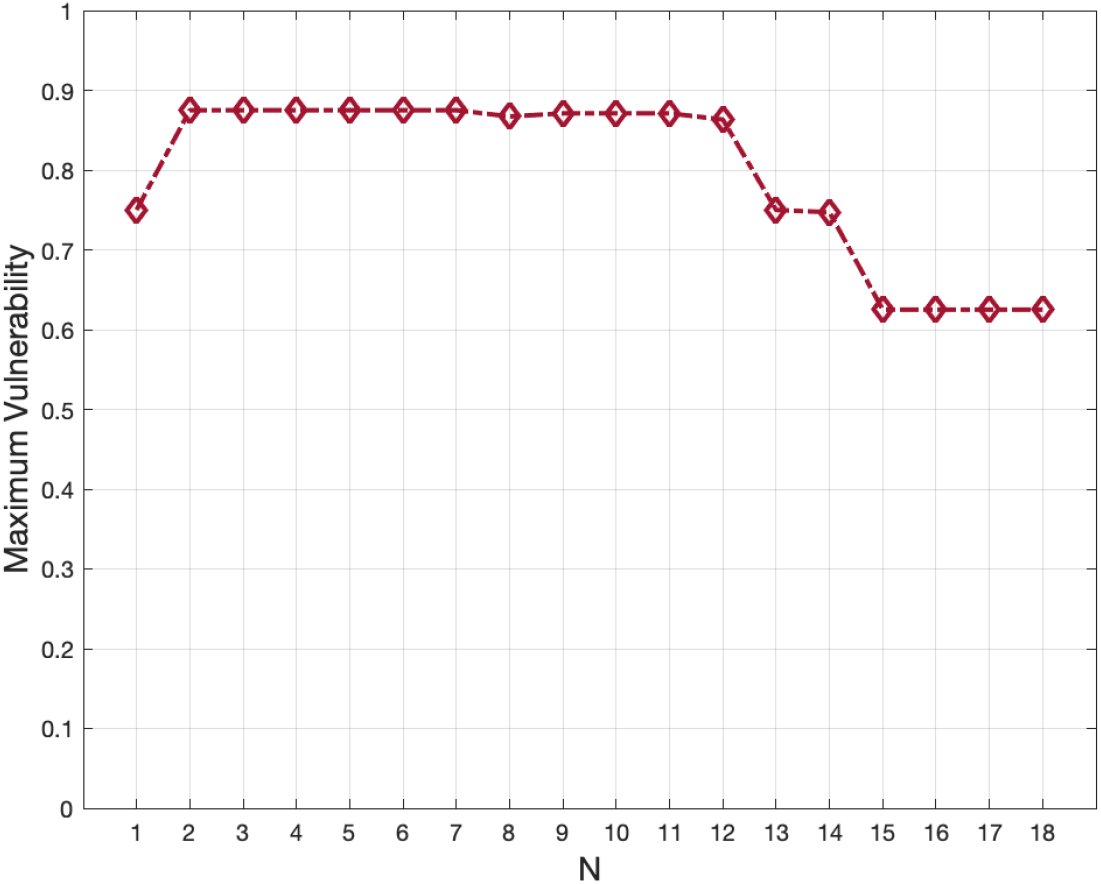
The ERBB signaling network maximum vulnerability levels, when there are *N* dysfunctional molecules in the network, computed using the proposed algorithm to study worst possible signaling failures.

An unexpected observation is that as the number of faulty molecules *N* increases, the maximum vulnerability values do not increase accordingly (Figure 2). While we see a maximum vulnerability increase going from single faults to double faults, i.e., *N* = 1 and 2 respectively, the maximum vulnerability quickly reaches a plateau and no longer increase further afterwards. Another interesting observation is that the smallest *N* for which we see the highest maximum vulnerability in this network is *N* = 2, i.e., double faults. This means there are some pairs of faulty molecules that cause the most detrimental damages to the signaling network, and increasing the number of faulty molecules can no further deteriorate the function of outputs.

### B. Extreme Failure Study of T cell Signaling Network

The T cell network (Figure 3) has three inputs and fourteen outputs. The input molecules are cd28 (cluster of differentiation 28), cd4 (cluster of differentiation 4) and tcrlig (ligand-bound T-cell receptor) [2]. For the output molecules we have shp2 (Src homology region 2 domain-containing phosphatase-2), bclxl (B-cell lymphoma-extra large), p70s, ap1 (activator protein 1), sre (serum response element), bcat (branched-chain amino acid transaminase), cyc1 (cytochrome c1), p21c, p27k, fkhr (forkhead transcription factor Foxo1), p38, cre (cAMP, cyclic adenosine monophosphate, response elements), nfat (nuclear factor of activated T-cells), and nfkb (nuclear factor kappa-light-chain-enhancer of activated B cells) [2]. Equations that specify how the activity of each molecule is regulated by its inputs are listed in Table 2.

**Figure 3.**
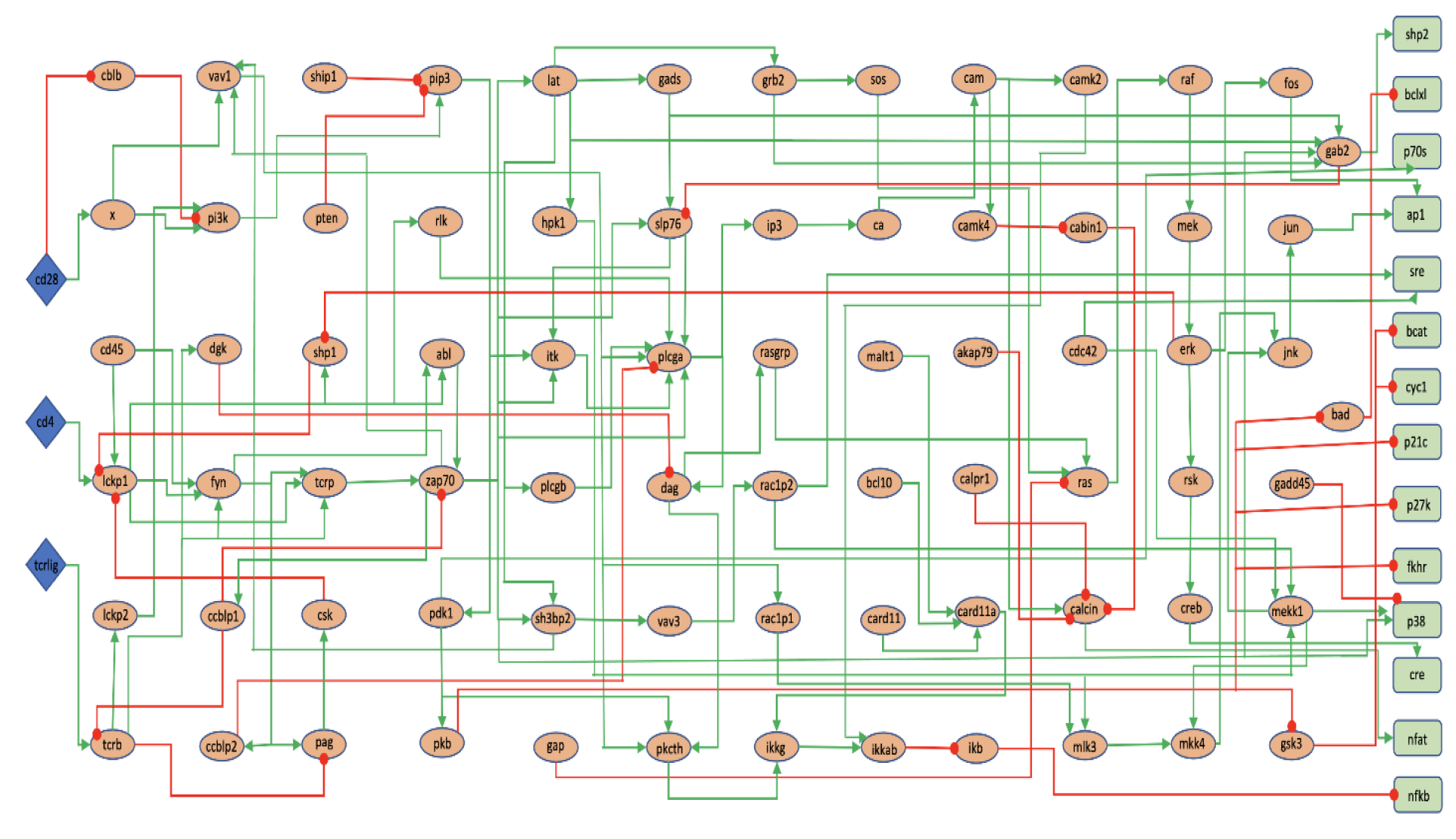
The experimentally-verified T cell signaling network reproduced from the study of Saez-Rodriguez et al [2]. The green arrows are activatory interactions and the red circle-ended edges are inhibitory interactions. The input nodes represent cd28, cd4, and tcrlig, whereas the output nodes stand for shp2, bclxl, p70s, ap1, sre, bcat, cyc1, p21c, p27k, fkhr, p38, cre, nfat and nfkb (see the main text for the acronyms and abbreviations).

**Table 2.**
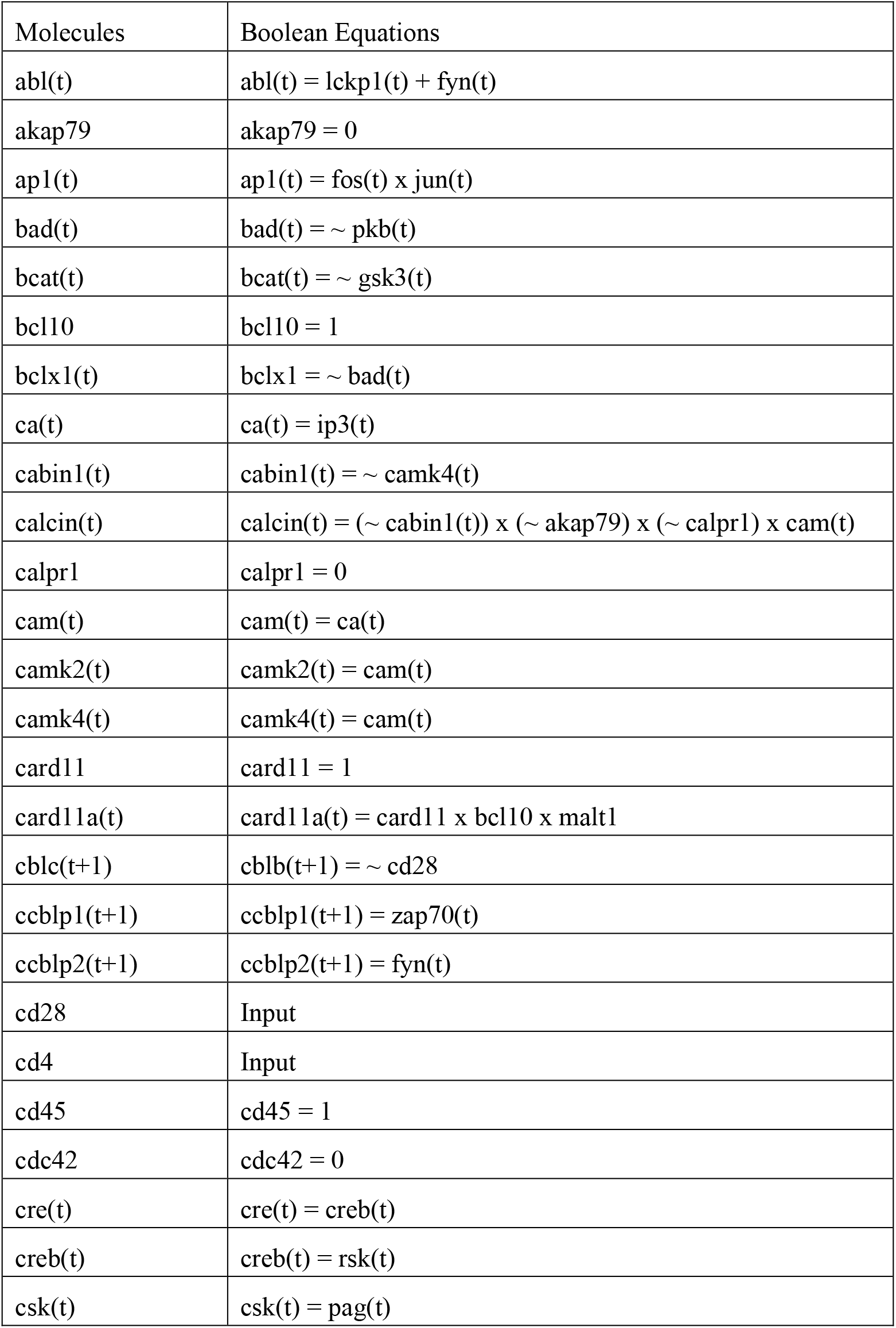
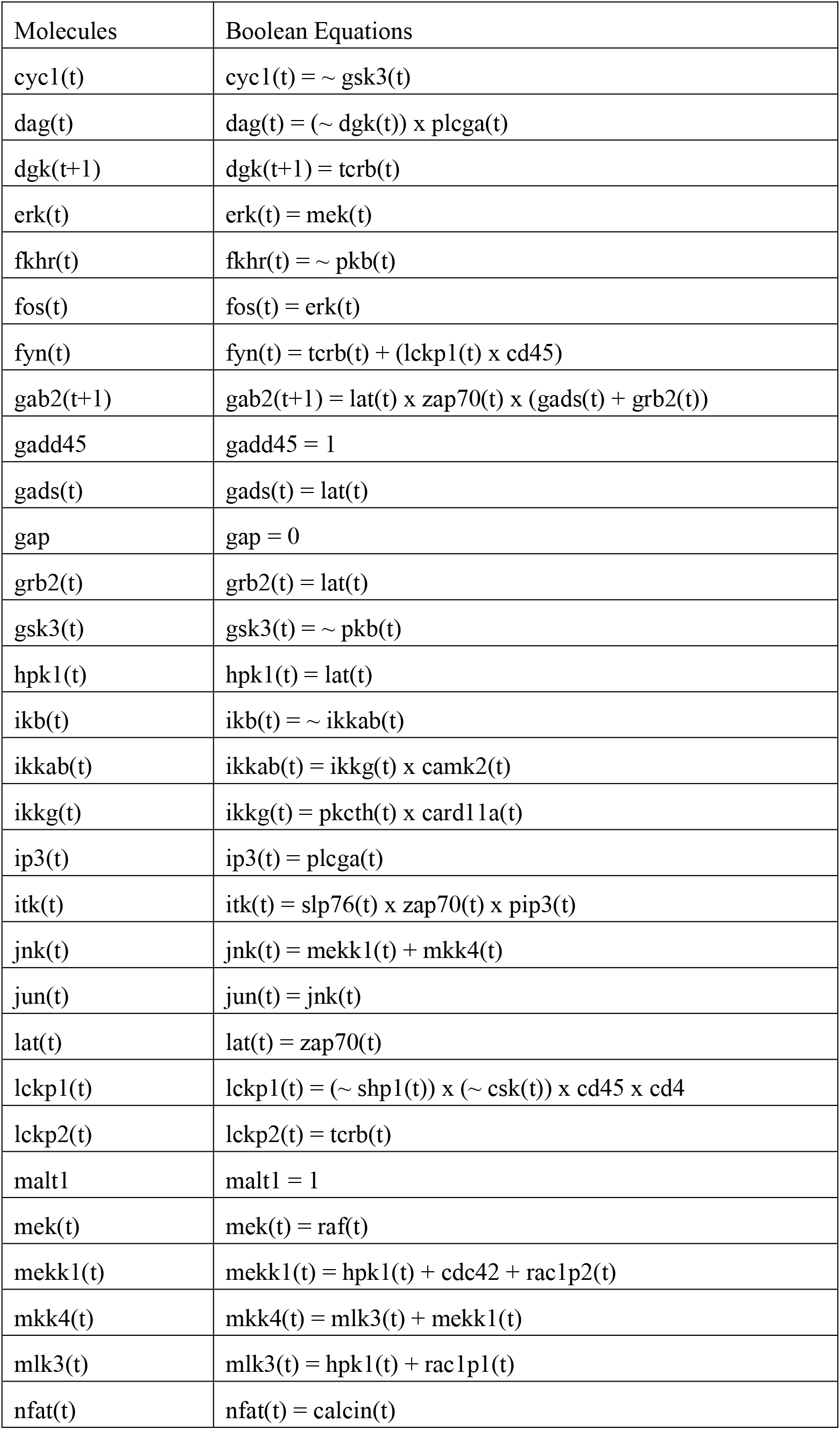
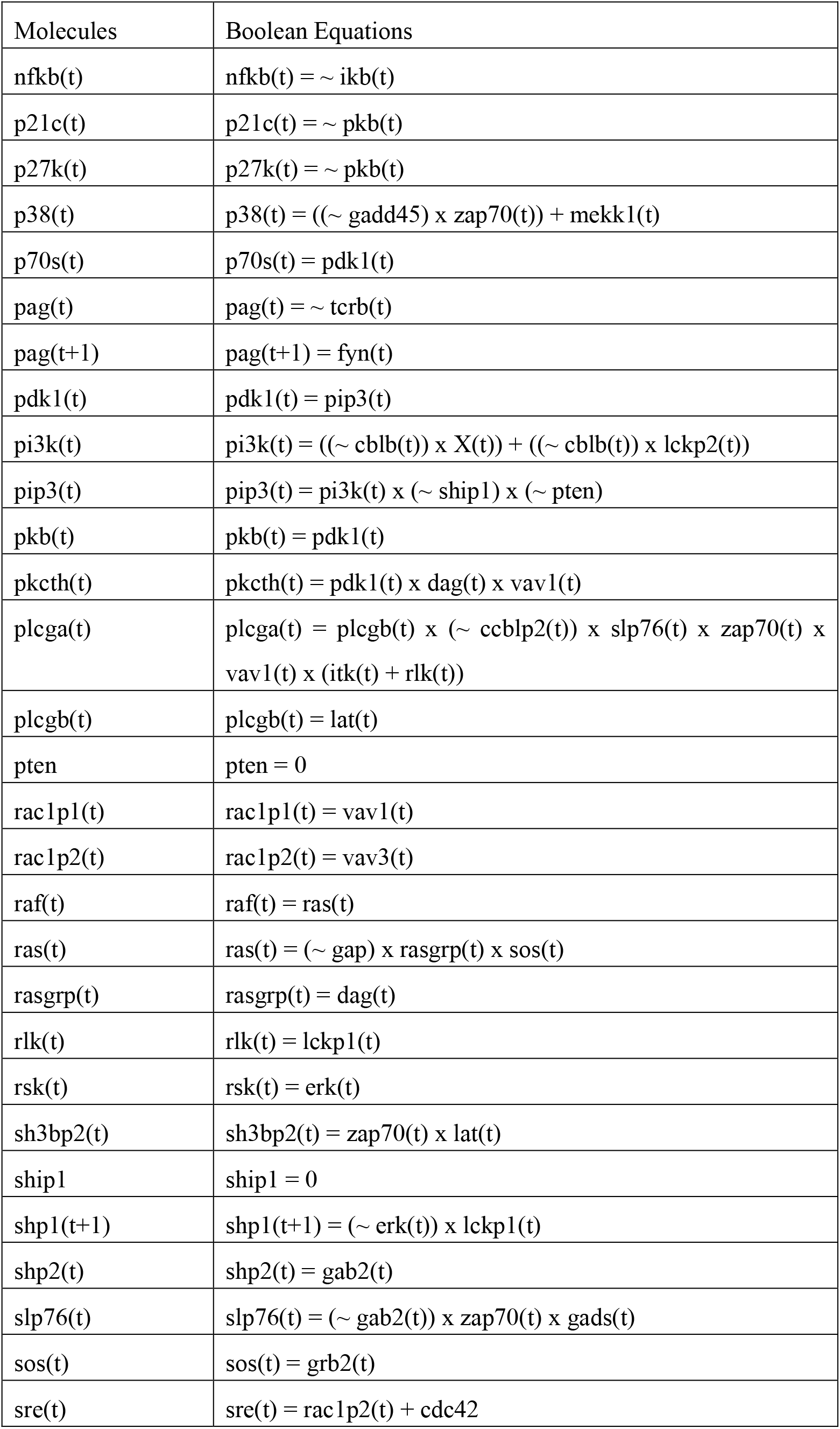
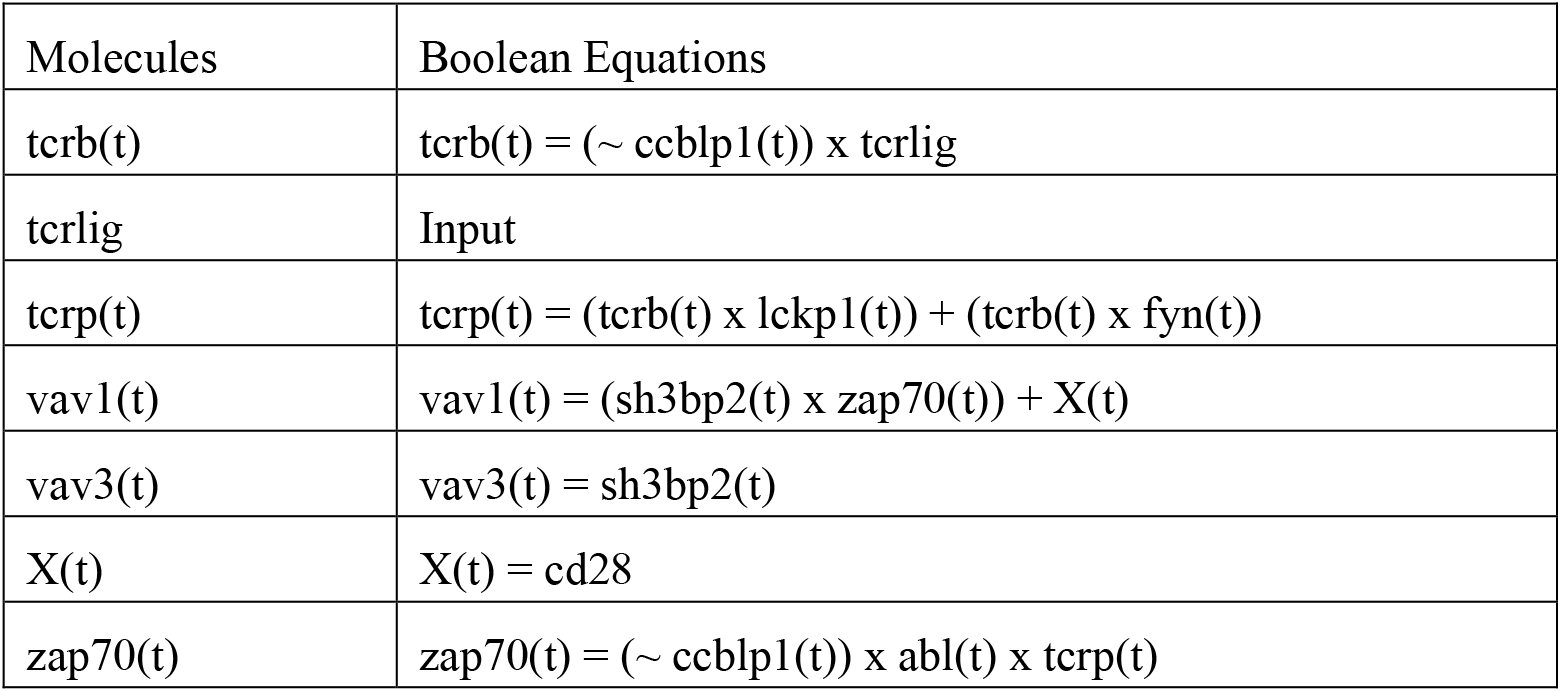
The Boolean equations for the T cell signaling network (Figure 3), provided in [2]. In the equations, “x” is used for the AND operation. “+” is used for the OR operation, and “∼” is used for the NOT operation. The symbol “t” represents the current time whereas “t+1” stands for the next time instant.

Using the developed vulnerability analysis equations (Methods, Section A) and the proposed extreme signaling failure analysis algorithm (Methods, Section B), the network maximum vulnerability is computed for each *N* (Figure 4), where *N* is the number of molecules that are simultaneously faulty. For any *N*, the network maximum vulnerability is the highest probability of network failure.

**Figure 4.**
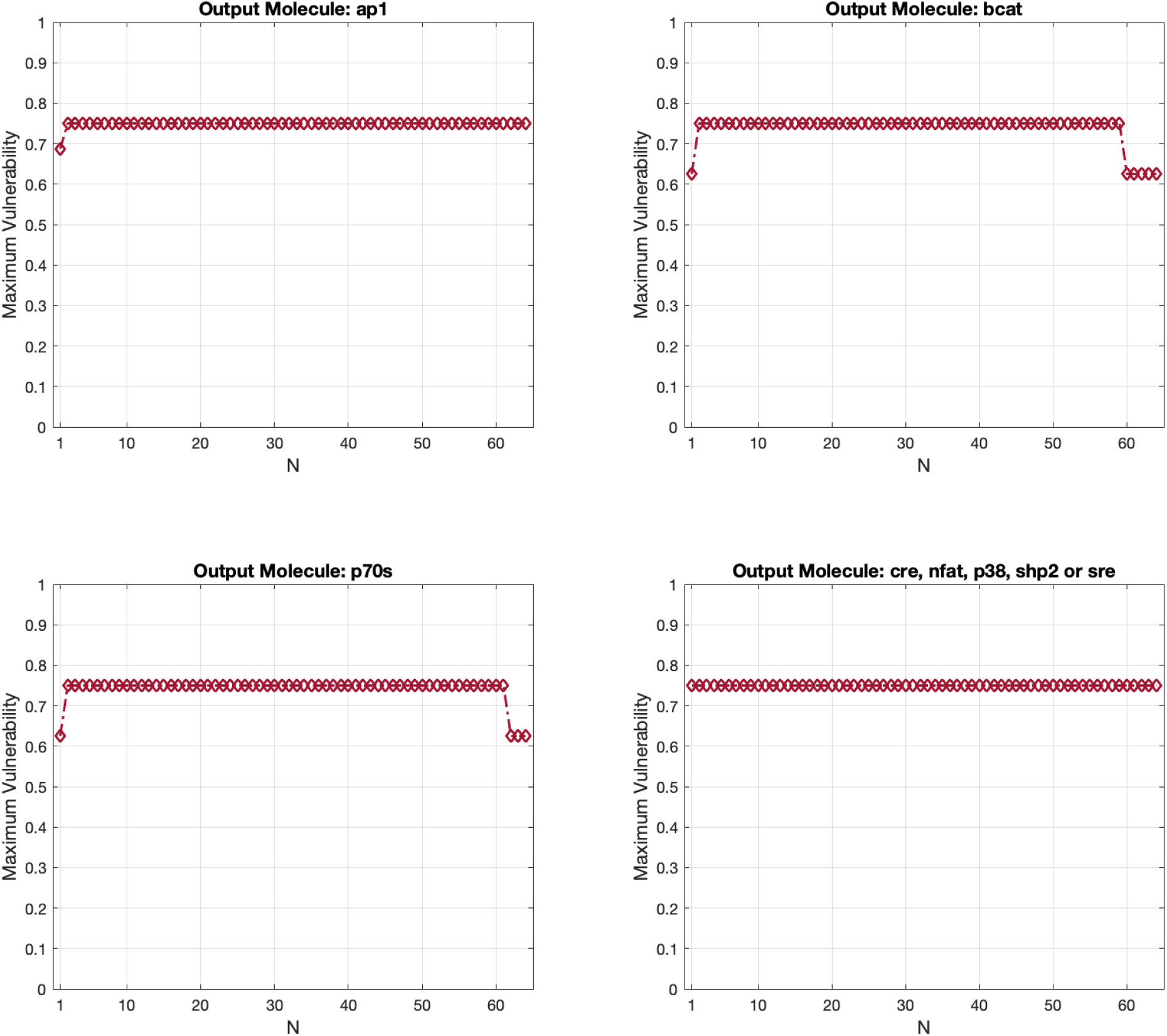
The T cell signaling network maximum vulnerability levels for the network outputs ap1, bcat, cre, nfat, p38, p70s, shp2 and sre, when there are *N* dysfunctional molecules in the network. The results are computed using the proposed algorithm to study the extreme signaling failures.

Similar to the ERBB network, but this time for all the outputs, we see as the number of faulty molecules *N* increases, again maximum vulnerability values do not increase accordingly (Figure 4). Moreover, for the network outputs ap1, bcat and p70s, while we see a maximum vulnerability increase going from single faults to double faults, *N* =1 and 2, respectively, the maximum vulnerability does not increase more afterwards. For these outputs, the smallest *N* for which we see the highest maximum vulnerability in this network is *N* = 2, i.e., double faults. This means that there are some pairs of faulty molecules that cause the most detrimental damage for these outputs, and increasing the number of faulty molecules does not further deteriorate the function of these outputs. However, for some other network outputs, including cre, nfat, p38, shp2 and sre, the results are different. For these outputs, the highest maximum vulnerability occurs when *N* = 1, which implies that there are some single faulty molecules that cause the can individually create the extreme signaling failures at these specific outputs (Figure 4).

### COMPUTATIONAL COMPLEXITY OF THE EXTREME SIGNALING FAILURE ALGORITHM

In this section, we determine the computational complexity of the proposed algorithm and compare it with the running time of exhaustive search. The extreme signaling failure analysis can be performed via an exhaustive search. This means that if it is of interest to find the maximum network vulnerability when there are *N* faulty molecules in the network, all possible groups of *N* faulty molecules have to be considered one by one, and the vulnerability value for each group needs to be individually computed. For example, consider a network with *K* = 50 molecules. When *N* = 2, the total number of *pairs* of faulty molecules that the exhaustive search has to examine can be shown to be 1,225 (see Equation (1) below). For *N* = 5, however, the total number of groups of *five* faulty molecules that the exhaustive search needs to consider increases to 2,118,760. This computational complexity becomes highly prohibitive as the network size *K* increases. In what follows, we show that the proposed extreme signaling failure algorithm is much less complex than the exhaustive search.

To determine and compare the computational complexities, let *K* be the number of molecules in a network, and *N* be the number of molecules that are simultaneously faulty in the network. The total number of groups of *N* faulty molecules out of *K* molecules, 1≤ *N* ≤ *K*, is given by the following equation, where *C*(*K, N*) represents the number of possible combinations

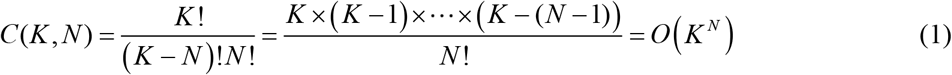

Here the *O*-notation stands for the asymptotic upper bound [20], with *K* being large. The term *O*(*K* ^*N*^) represents the computational complexity of *C*(*K, N*) as a function of *K* and *N*.

The computational complexity of the exhaustive search *α* (*K*) is the overall number of all possible groups of *N* faulty molecules for which vulnerabilities have to be computed, *N* = 1, …, *K*, i.e., *α*(*K*) = *C*(*K*,1) +… + *C*(*K, K*). To simplify the notation and without loss of generality, assume *K* is even. We note that *C*(*K, N*) has a maximum at *N* = *K* /2 [20], it is symmetric, i.e., *C*(*K, N*) = *C*(*K, K* − *N*), *N* = 1,…, (*K* / 2) −1, and *C*(*K, K*) =1. Therefore, the computational complexity of the exhaustive search simplifies to

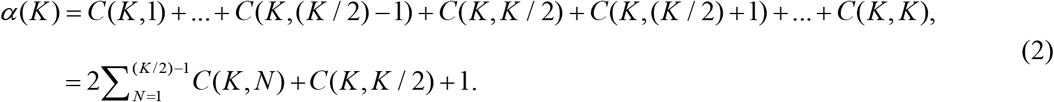

With *C*(*K,K*/2) being the dominant term in Equation (2) when *K* is large, and also using Equation (1), the exhaustive search computational complexity can be finally written as

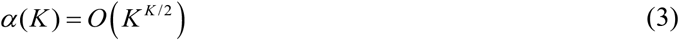

To determine the computational complexity of the proposed extreme signaling failure algorithm (Methods, Section B), we note that initially all groups of one, two and three faulty molecules are considered, *N* = 1, 2,3. And for the rest, *N* = 4,…, *K*, only *K* − (*N* − 1) vulnerabilities are computed. This is inspired by the observation [11] that typically a molecule with a high vulnerability appears in larger groups of molecules with high vulnerabilities, and based our experiments, *N* ≥ 4 is large enough and provides good accuracy, as discussed in the paragraph immediately after Equation (5). Therefore, the computational complexity *β* (*K*) of the algorithm can be written as

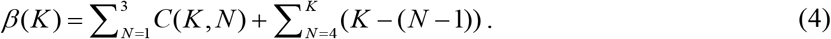

It can be verified that *C* (*K*, 3) in the above expression is the dominant term, when *K* is large. This simplifies the proposed algorithm computational complexity to

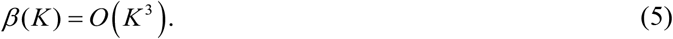

Upon comparing Equations (3) and (5), we note that since *O* (*K*^3^) ≪ *O* (*K K*/2), the proposed algorithm is much simpler and therefore much faster than the exhaustive search. For example, for a network with *K* = 50 molecules, the proposed algorithm complexity is in the order of 50^3^ ≈ 1.3 ×10^5^, which is much smaller than 50^25^ ≈ 3 ×10^42^, the exhaustive search complexity. With regard to the accuracy, we have observed that the differences between the results of the algorithm and the exhaustive search are 0.5% and 0%, for the ERBB and T cell networks, respectively, over all *N* values for which it was practical to perform the exhaustive search.

To practically compare the execution times of the proposed algorithm and the exhaustive search, we ran them on a computer with Intel Core i7 CPU, 3.4 GHz and 32 GB RAM. For the small ERBB signaling network (Figure 1), the exhaustive search took about 10 days for *N* = 1,…,18. In contrast and again for *N* = 1,…,18 (Figure 2), the proposed algorithm took only about 20 seconds, with an accuracy of 99.5%, compared to the exhaustive search results.

## METHODS

### A. Equations for Computing Vulnerability Levels

Computing the vulnerability level of a molecule or a group of molecules in a network can help to identify and to rank the key components of the network, and to discover appropriate therapeutics targets. Vulnerability of a molecule can be defined as the probability of having incorrect network responses when the molecule is dysfunctional. The dysfunction state of a molecule can be defined as failure to respond correctly to its input signals. In this paper, we consider stuck-at-0 (SA0) and stuck-at-1 (SA1) fault models to model the dysfunction of molecules, which are constantly 0, inactive, or 1, active, regardless of the activity state of the input signals of each molecule.

To compute the vulnerability level of a molecule, one needs to first introduce the sample space associated with the correct and incorrect network responses at the network output. Suppose *K* is the number of intermediate molecules in the network. Let *N* be the number of molecules that are simultaneously faulty, i.e., dysfunctional, *I* be the number of the network input combinations, CC be the number of clock cycles (time points) for which the network response is computed, and finally, let *l* be the subscript ranging from 1 to *C*(*K, N*), indexing faulty molecules or groups of faulty molecules (an algorithm for determining the required CC is given in Methods, Section C). Then, for each input combination stimulating the network in the presence of *N* faulty molecules, there will be CC number of output responses that may be <u>c</u>orrect, *c*, or <u>e</u>rroneous, *e*, in each clock cycle. Therefore, the sample space *S* can be defined as the set of all possible output sequences of *c* and *e* responses over CC clock cycles, that is

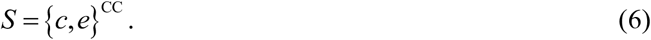

Moreover, for the *l*-th faulty molecule or the *l*-th group of faulty molecules, we define the event *S*_*l*_, a subset of *S*, as the set of all output sequences of *c* and *e* responses over CC clock cycles for all input combinations, that is

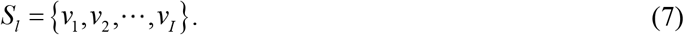

Note that *v*_*i*_ ∈ *S, i* =1, …, *I*, is a sequence of *c* and *e* of length CC, in which having an *e* in the *t*-th element of *v*_*i*_ means that an erroneous response is observed in the *t*-th clock cycle. Depending on the possible network responses and that which molecule or group of molecules is faulty, *v*_*i*_ s may have the same or different probabilities. Also note that some *v*_*i*_ s may be identical, therefore, *I* is indeed the maximum number of elements of *S*_*l*_. For CC = 2 and *I* = 2, for example, we have *S*_*l*_ ={*v*_1_, *v*_2_}, where *v*_1_,*v*_2_ ∈ *S* ={*c, e*}^2^ ={(*c, c*),(*c, e*),(*e, c*),(*e, e*)}.

In addition, we define CC number of events as follows: *E*_1_ = the event of having an erroneous network response at the output in the 1^st^ clock cycle, …, and *E*_CC_ = the event of having an erroneous network response at the output in the CC^th^ clock cycle. Note that *E*_1_ is the set of those *v* elements in *S*_*l*_ in Equation (7) that have an *e* as the 1^st^ entry, *E*_2_ is the set of those *v* elements in *S*_*l*_ that have an *e* as the 2^nd^ entry, and so on. We define the vulnerability level of the *l*-th molecule *M*_*l*_, Vul(*M*_*l*_), as the probability of having an erroneous network response in the 1^st^ clock cycle, …, or in the CC^th^ clock cycle, when dysfunctional. Therefore, Vul(*M*_*l*_) can be written as

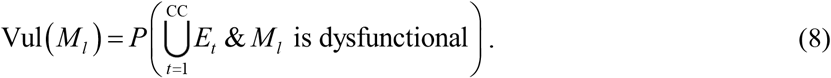

Since we consider SA0 and SA1 as the fault models, Equation (8) can be expanded as follows

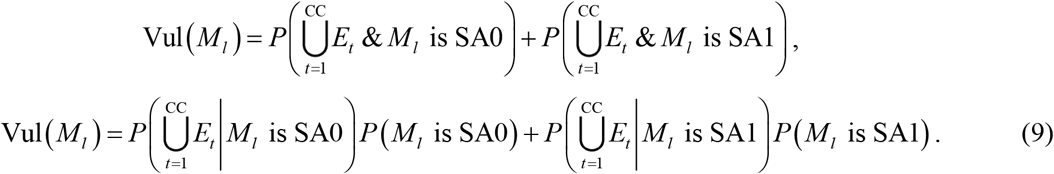

While we assume equi-probable SA0 and SA1 faults for each molecule, i.e., *P* (*M*_*l*_ is SA0) = *P* (*M*_*l*_ is SA1) = 0.5 in our computations, Equations (8) and (9) can be extended to other fault models and fault probabilities.

Equation (9) is provided for computing the vulnerability level of a single molecule. However, it is also of interest to study the abnormal network responses when multiple molecules are faulty at the same time. In the large T cell network (Figure 3) and for simplicity, in the extreme signaling failure analysis (Figure 4), we assume *N* molecules are either all SA0 or all SA1 at the same time, *N* > 1. For this scenario, Equation (9) is still applicable, where *M*_*l*_ needs to be merely replaced with “ *M*_1_, *M* _2_, …, *M*_*N*_ “. On the other hand, for the small ERBB network (Figure 1) and its extreme signaling failure analysis (Figure 2), we consider all possible SA0 and SA1 fault combinations for *N* dysfunctional molecules, *N* > 1. In this scenario, we extend Equation (9) as follows

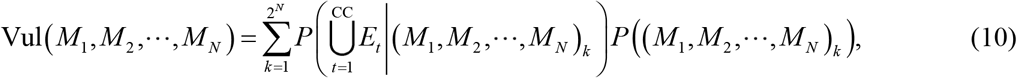

where (*M*_1_, *M*_2_, …, *M*_*N*_)_*k*_ ∈{SA0, SA1}^*N*^ is the *k*-th fault vector for the group of *N* dysfunctional molecules *M*_1_, *M* _2_, …, and *M*_*N*_.

**Example:** In the ERBB signaling network (Figure 1) we have *K* =18, *I* = 2, and CC = 5, with the CC being determined using the algorithm introduced and developed in Methods, Section C. When *N* = 1, a single faulty molecule, there are two events associated with the *l*-th faulty molecule *M*_*l*_ being SA0 or SA1

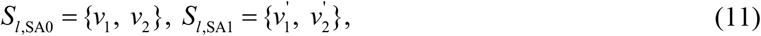

where 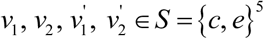 for all *l* = 1, …, 18. Using Equation (9), single-fault vulnerability levels of molecules are calculated for all *l* = 1, …, 18. To illustrate and for CC = 2, assume hypothetically that for the *M*_*l*_ faulty molecule we have *S*_*l*, SA0_ ={(*c, c*), (*e, e*)} with the probabilities of *P*(*v*_1_) = 0.5 and *P*(*v*_2_) = 0.5, and *S*_*l*, SA1_ ={(*c, e*)} with the probability of 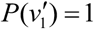. Based on the definition of *E*_*t*_, it can be shown that *E*_1_ = *E*_2_ ={(*e, e*)} when *M*_*l*_ is SA0, whereas *E*_1_ =Ø and *E*_2_ = {(*c, e*)} when *M*_*l*_ is SA1. Using Equation (9), vulnerability level of *l*-th faulty molecule can be computed for equi-probable SA0 and SA1 faults as follows

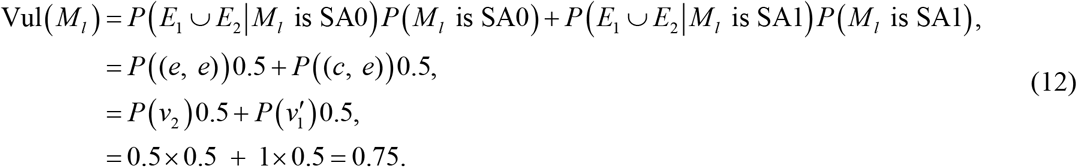

Similarly, when of faulty molecules *N* = 2, a pair of faulty molecules, there are four events associated with each pair of faulty molecules

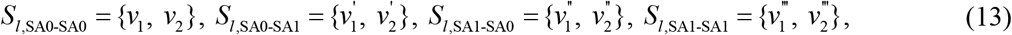

where 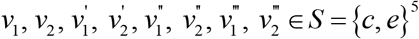, for all *l* = 1, …, 153 (there are 153 possible pairs, i.e., *C* (18, 2) = 153). Then, Equation (10) is used to compute the double-fault vulnerability level of each pair as follows

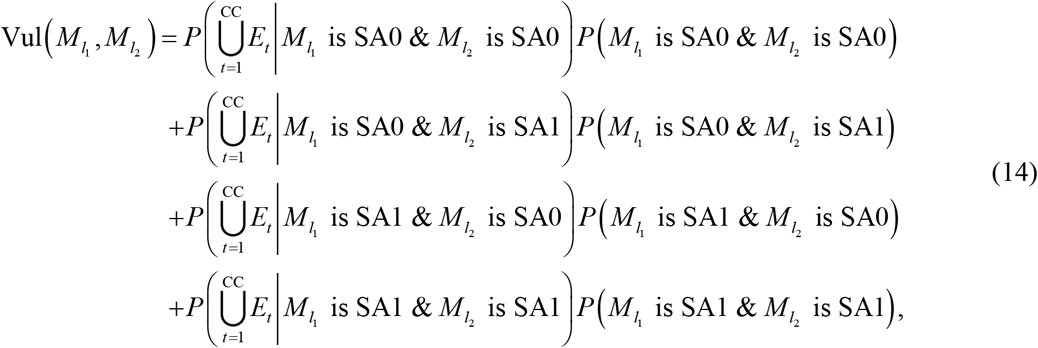

Where *l*_1_, *l*_2_ = 1, …,18, *l*_1_ ≠ *l*_2_, with equi-probable and independent SA0 and SA1 faults. These steps are repeated for all *N* > 2, to complete the ERBB network extreme signaling failure analysis. Note that what we observe in Figure 2 are maximum vulnerabilities, e.g., max_*l*_ Vul(*M*_*l*_) and 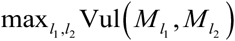 for *N* =1 and 2, respectively. To compute the vulnerability levels for the T cell signaling network, the considered parameter values are *K* = 64, *I* = 8, and CC = 5.

### B. Proposed Algorithm for the Extreme Signaling Failure Analysis

I. In this section, we provide a detailed explanation of the proposed algorithm. The extreme signaling failure analysis can be performed by an exhaustive search. However, the time needed by the exhaustive search grows exponentially as the size of the network increases, as presented earlier in the paper. To avoid this high computational complexity, we propose the main algorithm with the following four steps
II. First, we compute an upper bound on the number of clock cycles needed for computing the vulnerability levels (Methods, Section C), so that we prevent running network simulations longer than what is needed.
III. Next, we use Equation (10) to compute 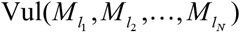 for *N* =1,2, and 3. This is motivated by the observation [11] that typically a molecule with high vulnerability appears in larger groups of molecules with high vulnerabilities, and based our experiments, *N* ≥ 4 is large enough and provides good accuracy, as described in the paragraph immediately after Equation (5). Thus far,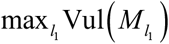, 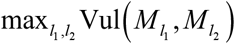 and 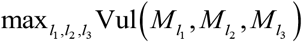 represent the extreme signaling failures when there are single, double and triple faults, respectively.
IV. To determine the extreme signaling failure when there are four simultaneously faulty molecules, *N* = 4, we pick the molecular triplet, group of *N* − 1 faulty molecules, having the highest vulnerability value, e.g., (*a,b, c*). Then we compute vulnerabilities only for those *K* − (*N* − 1) quadruplets, groups of *N* faulty molecules with *N* = 4, that include (*a,b, c*), i.e., (*a,b, c, M*_*l*_). This results in max_*l*_ Vul(*a,b, c, M*_*l*_) as the extreme signaling failure when there are *N* = 4 simultaneous faults.
V. Then, we repeat Step III for *N* = 5,…, *K*, to complete the extreme signaling failure analysis.

Note that this algorithm is not limited to a specific molecular network. Furthermore, in addition to the vulnerability parameter used in this paper, other parameters that quantify and rank the importance of a molecule or a group of molecules can be used in the algorithm as well.

### C. Proposed Algorithm for Determining the Number of Required Clock Cycles to Compute the Vulnerabilities

Modeling and analysis of molecular networks become more challenging if there are positive or negative feedback paths. Due to the feedback mechanisms, network responses may change over time because of some internal compensatory or regulatory mechanisms [18,19]. Feedbacks can cause delays in propagation of signals to the network outputs, while passing through the feedback paths. Therefore, analysis of the effects of feedback in computing vulnerability levels of the network molecules is of interest. More precisely, in this paper we are interested in determining how many clock cycles are needed to compute the vulnerability level of a molecule or a group of molecules, when there are feedbacks in the network. For this purpose, in this section we propose an algorithm that computes an upper bound on the number of clock cycles needed to generate the network response, to calculate its molecular vulnerabilities. Using this algorithm, one can specify how many times the network needs to be simulated, for a normal or abnormal signal to complete its propagation to the network output. This is needed in the proposed main algorithm for the extreme signaling failure analysis (Methods, Section B), to minimize the overall simulation time.

The feedback paths in a network can be modeled by unit-delay memory elements called flip-flops [9]. In a network, if there is only one feedback path, then we intuitively need at most two clock cycles to see full effect of an error, i.e., the effects of an incorrect signal value of a faulty molecule on possibly other molecules and pathways, that collectively determine the network output response. This is because of the delayed response of the flip-flop in the feedback path. In fact, if after the 1^st^ clock cycle there exists an erroneous signal value of a faulty molecule at the input of the feedback flip-flop, then the 2^nd^ clock cycle may be needed for that error to show its full effect at the network output. This is because the feedback-delayed erroneous signal of the faulty molecule may affect some other molecules and pathways in the 2^nd^ clock cycle, which may increase the probability of incorrect responses at the network output. In general, if there exist *F* feedback paths in the network, then we need to simulate the network for at most *F* +1 clock cycles, for an error to show its full effect at the network output. This maximum number of clock cycles is required, if these two conditions hold: (i) all the feedback paths are in the same pathway, connected in series and exhibiting *F* feedback flip-flops; and (ii) after the 1^st^ clock cycle, an error appears at the input of the first feedback flip-flop.

The upper bound of *F* +1 clock cycles can be tightened, by finding the pathways within the network containing the highest number of feedback paths in series. For instance, if there exist *F* feedback paths in the network, with *L* ≤ *F* being the largest number of feedback paths in series on the same pathway, then the maximum clock cycles needed is bounded above by *L* +1, i.e., the number of clock cycles needed for an error to show its full effect is less than or equal to *L* + 1. Therefore, to determine the maximum number of required clock cycles, it is sufficient to examine the connections between the network feedback paths and determine how many of them are serially connected in a single pathway. To do this, we define a graph theory topological metric called *closeness* (*CL*). The 0 ≤ *CL* (*X, Y*) ≤ 1 parameter for quantifying the closeness of two molecules *X* and *Y* in a network is the inverse of the distance *d* (*X, Y*) between the two molecules, defined as the length of the shortest path between the two molecules in the network graph

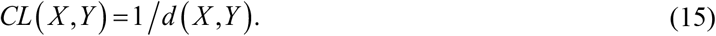

Some examples of how to calculate the closeness parameter are presented in Figure 5.

**Figure 5.**
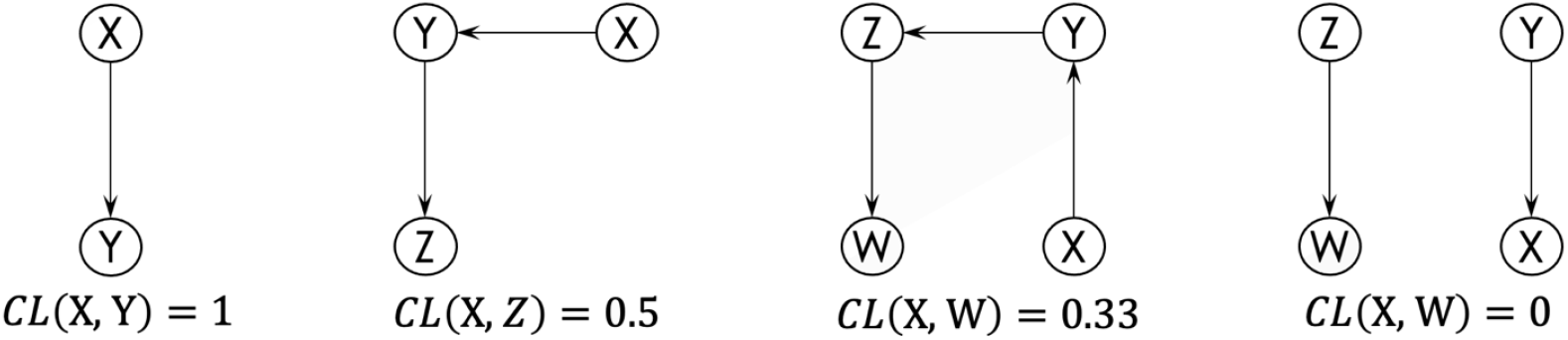
Numerical examples of the closeness parameter *CL* between two molecules.

To determine the maximum number of clock cycles needed for computing the vulnerability levels in a network, we propose the following algorithm with four steps

I. Identify the feedback paths and the nodes that initiate the feedbacks, in the network. If they are not known a priori, identify them by finding the loops in the network graph, using a graph algorithm such as the depth-first search algorithm [20].
II. Assume there exist *F* feedback paths in the network. Arbitrarily label the feedback initiating nodes as *f*_*i*_, *i* = 1,…, *F*. Then start with *i* = 1, by calculating the closeness of *f*_1_ with respect to the other feedback initiating nodes *f* _*j*_ s, *j* ≠ *I* and *j* =1,…, *F*. If *CL*(*f*_1_, *f* _*j*_) = 0 for all *j* values, it means *f*_1_ is on no other pathway with other feedback initiating nodes, and now *f*_2_ has to be examined similarly, i.e., *i* = 2, *j* ≠ *I* and *j* =1,…, *F*, and so forth. However, if *CL*(*f*_1_, *f* _*j*_) ≠ 0 for *j* = *j*_0_, then this indicates that there is a path between *f*_1_ and 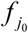, i.e., the feedback initiating nodes *f*_1_ and 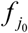 are on the same pathway. This finding needs to be pursued in the next step.
III. For fixed *i* and *j* values, e.g., *i* = 1 and *j* = *j*_0_, calculate *CL*(*f* _*j*_, *f*_*k*_) for all other feedback initiating nodes *f*_*k*_ s, *k* ≠ *i, j*, and *k* =1,, *F*. Depending on the *CL* being non-zero or zero, it can be identified if *f*_*k*_ is on the same pathway that includes both *f*_*i*_ and *f* _*j*_ feedback initiating nodes or not.
IV. Repeat Step III until all the feedback initiating nodes are examined.

Using the information obtained from executing the above steps, the algorithm finds the pathway that contains the largest number of feedback initiating nodes on it, in series. With this specific number being called *L*, then at most *L +* 1 clock cycles are needed for computing the vulnerability levels.

#### Toy examples

Here we compute vulnerability levels in two toy networks (Figure 6A and Figure 6C) that have different number of feedback paths, to describe the relation between *F* and vulnerabilities. Note that since each network has a single pathway, we have *L* = *F* in both networks.

**Figure 6.**
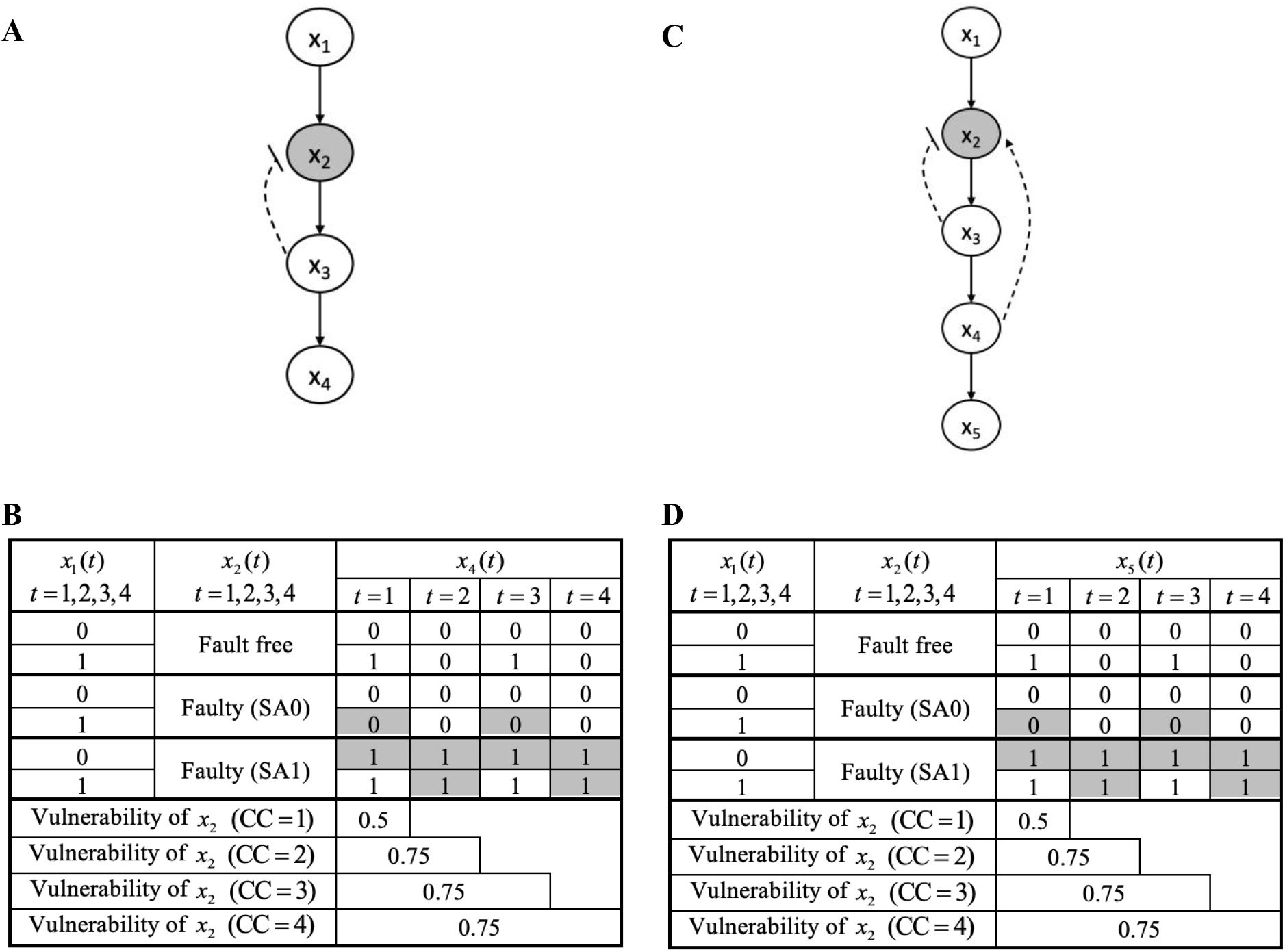
Toy networks illustrating the number of clock cycles needed for the erroneous signal of a faulty (dysfunctional) molecule to show its full effect at the network output. (A) Toy network with one feedback path. (B) Output truth table for fault-free and faulty *x*_2_. (C) Toy network with two feedback paths. (D) Output truth table for fault-free and faulty *x*_2_. Note: The dashed lines ending in an arrowhead or a blunt line represent positive and negative feedbacks, respectively.

The first toy network (Figure 6A) has four nodes: *x*_1_(*t*) is the input node (molecule), the intermediate nodes are *x*_2_ (*t*) = *x*_1_ (*t*) × (∼ *x*_3_ (*t* −1)) and *x*_3_ (*t*) = *x*_2_ (*t*), and *x*_4_ (*t*) = *x*_3_ (*t*) is the output node, where “ × “ is used for the AND operation and “ ∼ “ is used for the NOT operation, and *x*_3_ (0) = 0. Herein, the node *x*_3_ initiates a negative feedback to the node *x*_2_. Since there is only one feedback path in the network, *F* = 1, at most two clock cycles are enough, *F* +1 = 2, to observe the full error effect at the output. To demonstrate this, we compute the vulnerability level of the node *x*_2_ (Figure 6B), for different number of clock cycles CC = 1, 2,3, 4, using Equation (9) (Methods, Section A). We observe that the vulnerability of *x*_2_ with 1 clock cycle is 0.5 and it becomes 0.75 with 2 clock cycles, and then remains at 0.75 with 3 and 4 clock cycles. These indicate that the full vulnerability of *x*_2_ is 0.75 that is determined by analyzing the network having feedback for two clock cycles (*F* + 1 = 2). In other words, two clock cycles are needed for the erroneous *x*_2_ signal values to show their full effects at the network output *x*_4_. Additionally, more clock cycles are not needed.

The second toy network (Figure 6C) has five nodes: *x*_1_ (*t*) is the input node, *x*_2_ (*t*) = (*x*_1_ (*t*) + *x*_4_ (*t* −1)) ×(∼ *x*_3_ (*t* −1)), *x*_3_ (*t*) = *x*_2_ (*t*) and *x*_4_ (*t*) = *x*_3_ (*t*) are the intermediate nodes, and *x*_5_ (*t*) is the output node, where “ + “ is used for the OR operation, and *x*_3_ (0) = *x*_4_ (0) = 0. Here the nodes *x*_3_ and *x*_4_ initiate a negative feedback and a positive feedback to the node *x*_2_, respectively. Since there are two feedback paths in the network, *F* = 2, at most three clock cycles are needed, *F* +1 = 3, to observe the full error effect of *x*_2_ at the output. The computed vulnerability level of *x*_2_ (Figure 6D) for different number of clock cycles corroborates what we stated earlier in this section, that is, *F* +1 is indeed an upper bound and less number of clock cycles may be needed for computing vulnerabilities in a network with feedbacks. In fact, we observe that the full vulnerability of *x*_2_ is 0.75, obtained using 2 clock cycles only, and analyzing the network for the upper bound of *F* +1 = 3 clock cycles is not needed (Figure 6D). In other words, 2 clock cycles are enough for errors originated from *x*_2_ to show their complete effects at the output *x*_5_.

### D. ERBB Signaling Network – Vulnerability and the Number of Clock Cycles

In this section, we apply the proposed algorithm in Methods Section C, to the ERBB signaling network (Figure 1). We start by identifying feedbacks in the network. Given the small size of the network, visual inspection of the network reveals five loops, which topologically may contain the indicators of feedback interactions. The five loops are (i) IGF1R →AKT1 →IGF1R, (ii) IGF1R →MEK1 →ER-*α* →IGF1R, (iii) CDK4 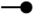 p27 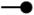 CDK4, (iv) CDK4 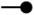 p21 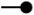 CDK4, and (v) CDK2 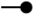 p27 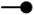 CDK2. Despite the existence of five loops, there are only four feedback initiating nodes, AKT1, ER-*α*, p21 and p27, because p27 is common between two loops, i.e., the loops (iii) and (iv). Note that in the absence of prior information about the feedbacks, the feedback initiators may not be uniquely determined within the identified loops. For example, based on these five loops, one can identify IGF1R, MEK1, CDK4 and CDK2 as feedback initiators as well. Nevertheless, different choices for the feedback initiating molecules within these loops do not affect the algorithm that aims at finding the pathway that contains the largest number of feedback initiating nodes on it, in series. This is because if the feedback initiating nodes *f*_*i*_ and *f* _*j*_ chosen from two loops are connected through a pathway, i.e., *CL*(*f*_*i*_, *f* _*j*_) ≠ 0, then other choices of the feedback initiating nodes 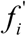 and 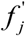 from the said two loops will be connected through a pathway as well, i.e., 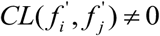. Thus, the algorithm to determine *L* is independent of the choice of the feedback initiating nodes.

Using the identified feedback initiators, the algorithm outputs the upper bound of *L* + 1 = 5 clock cycles that may be needed for computing the vulnerability level of a molecule. This is because the algorithm identifies a pathway that contains all the feedback initiating molecules on it, in series. For instance, AKT1 →ER-*α* →p27 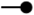 CDK4 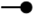 p21 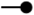 CDK2 →pRB is a pathway that contains all of the feedback initiating molecules on it. Through this specific pathway, erroneous signals originated from an upstream molecule of AKT1 may require five clock cycles to show their full effects on the output molecule pRB, due to the signal propagation delays introduced by the feedback paths connected in series on the same pathway. Computed using Equation (9) (Methods, Section A), Figure 7 presents the single-fault vulnerability levels versus the number of clock cycles CC for some molecules in the ERBB signaling network. We observe that the vulnerability levels of the molecules can be computed in less than five clock cycles, which confirms that it is sufficient to simulate and analyze the network for at most five clock cycles, as specified by the algorithm.

**Figure 7.**
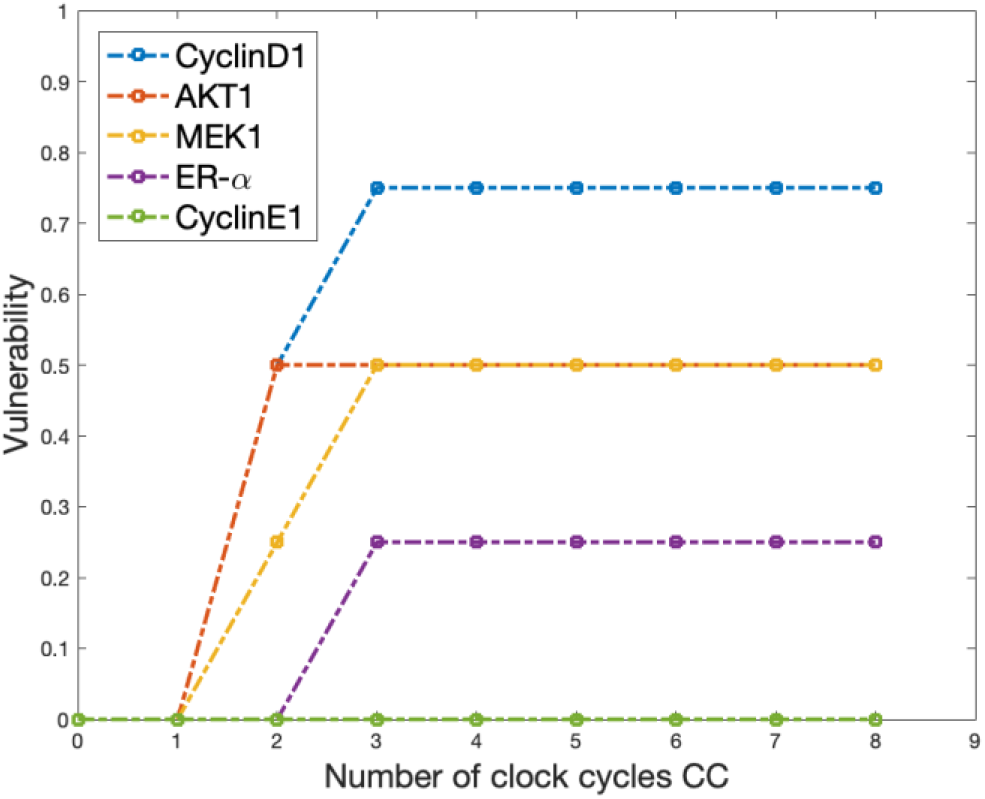
Vulnerability versus the number of clock cycles CC for some molecules in the ERBB signaling network.

### E. T Cell Signaling Network – Vulnerability and the Number of Clock Cycles

In this section, we apply the proposed algorithm in Methods, Section C to the T cell signaling network (Figure 3). In the network, first we identify four feedback initiating nodes that are shp1, ccblp1, pag, and gab2, using the time indices in Table 2 and also by visual inspection of Figure 3. After using the proposed algorithm, we obtain the upper bound of *L* + 1 = 5 clock cycles, because there is one pathway that contains all the four feedback initiating nodes in series. Next, we compute the single-fault vulnerability levels of some molecules versus the number of clock cycles CC (Figure 8A), with cre considered as the output molecule, and using Equation (9) (Methods, Section A). We observe that the vulnerability levels of the molecules can be computed in less than five clock cycles, which confirms that it is sufficient to simulate and analyze the network for at most five clock cycles, as determined by the algorithm.

**Figure 8.**
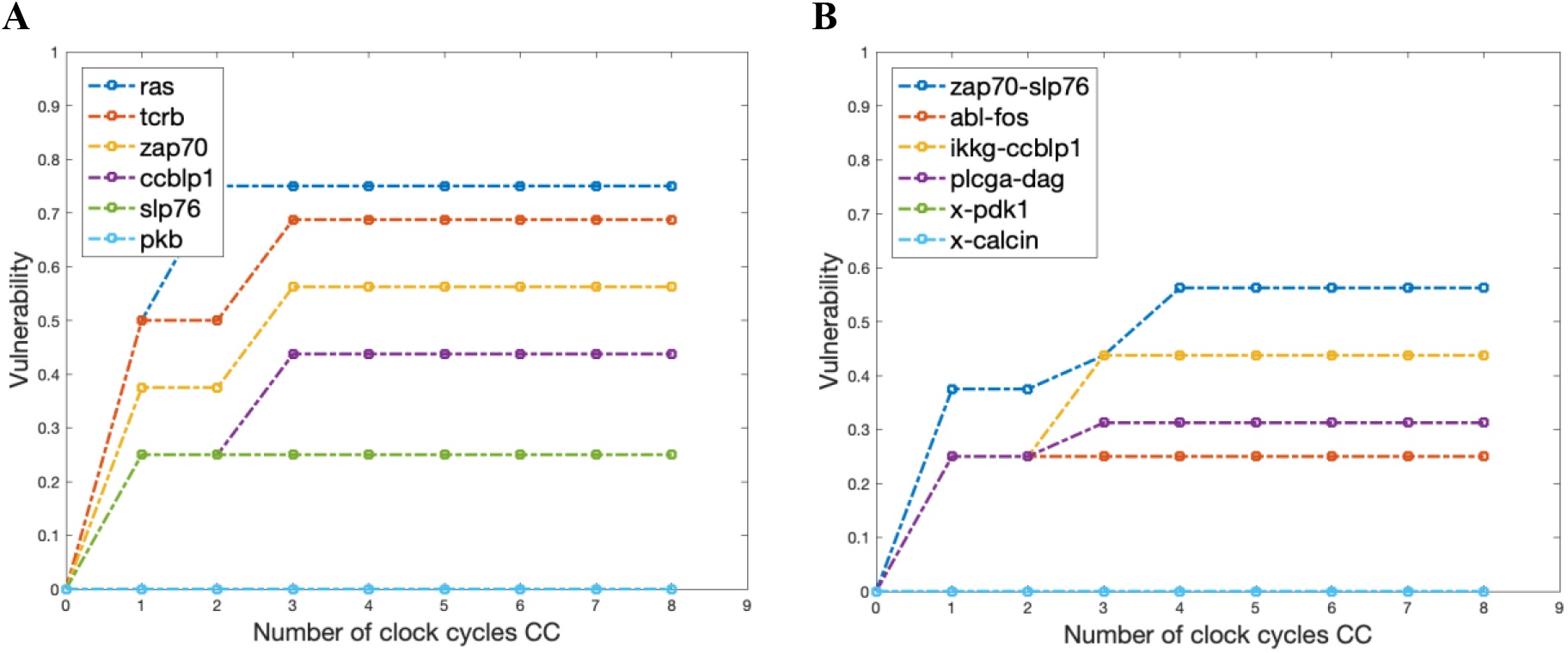
Vulnerability versus the number of clock cycles CC for some single and pairs of molecules in the T cell signaling network, with “cre” considered as the output molecule. (A) Single-fault vulnerability levels. (B) Double-fault vulnerability levels.

Furthermore, we compute the double-fault vulnerability values of some pairs of molecules versus the number of clock cycles CC (Figure 8B), with cre considered as the output molecule, and using Equation (9) (Methods, Section A). As anticipated by the algorithm, the vulnerability levels of the molecular pairs can be computed in less than five clock cycles, i.e., it is enough to simulate and analyze the network for at most five clock cycles, irrespective of considering single faults or double faults.

A noteworthy point is that when two molecules are faulty at the same time, more clock cycles may be needed to observe the aggregated full effects of the two erroneous signals at the network output, compared to single faults. However, the upper bound found by the algorithm still works for both scenarios. This is because the upper bound depends only on the topological positions of the feedback initiating nodes and the connections among them, and not on how many nodes are grouped together, to represent a group of faulty molecules.

As some numerical examples, consider the scenario of zap70 and slp76 being individually faulty (Figure 8A), where three and one clock cycles are needed, respectively, to compute their full single vulnerabilities of 0.56 and 0.25, respectively. On the other hand, when they are simultaneously faulty (Figure 8B), four, i.e., more clock cycles are needed to compute their full double vulnerability of 0.56. This indicates that if multiple molecules are faulty concurrently, then there may be further delays in observing the entire effects of multiple erroneous signals, propagating via various pathways towards the network output. Additionally, we observe that the upper bound of *L* + 1 = 5 clock cycles holds true for computing both single and double vulnerabilities.

## CONCLUSION

Signaling networks in cells are involved in controlling various cellular functions, through different signaling processes. Developing novel methods for the functional analysis of signaling networks is particularly important for understanding such complex normal and abnormal signaling processes, and can help for untangling the pathology of complex trait diseases. A fundamental question in systems biology is how vulnerable a signaling network is to the dysfunction of an individual or groups of molecules, where the dysfunction of a molecule is defined as failure to respond correctly to the input signals. The vulnerability levels associated with the dysfunction of molecules or groups of molecules can be measured by computing probabilities of having incorrect network responses in the presence of dysfunctional molecules, using the equations introduced and developed in Methods, Section A.

The focus of this study is to understand that, in a given network, what molecule or group of molecules can result in the most detrimental failure in the function of the network. To answer this question, here we propose a systematic method to identify extreme signaling failures in molecular networks (Methods, Section B). The extreme signaling failure here is defined to describe a pathological phenomenon in which the failure of the network passes the physiological tolerable noise threshold, which is quantified here as the maximum vulnerability level. The said pathological phenomenon is characterized to be emerged from the presence of one or more dysfunctional molecules in the network. While it is conceivable that different individual dysfunctional molecules may have different vulnerability levels, it is not clear what happens to the vulnerability levels, if two or more molecules are dysfunctional simultaneously.

The extreme signaling failure analysis is initially conducted on the ERBB signaling network (Figure 1). We observe that the maximum vulnerability values do not increase accordingly as the number of concurrently faulty molecules *N* increases (Figure 2). More precisely, we see a maximum vulnerability increase going from single faults to double faults, *N* = 1 and 2, respectively, and then the maximum vulnerability no longer increases when *N* increases. Moreover, we observe that the smallest *N* for which we see the highest maximum vulnerability in this network is *N* = 2, i.e., the double faults. This observation shows that in a network of 18 intermediate molecules, some selective pairs of faulty molecules are sufficient to create the most detrimental harm to the function of the output molecule, and increasing the number of simultaneously faulty molecules does not induce a worse scenario compared to having some specific dysfunctional pairs of molecules.

When extreme signaling failure analysis was done on the T cell signaling network (Figure 3), similar to the ERBB network results, we notice that the maximum vulnerability values do not increase accordingly as the number of simultaneously faulty molecules *N* increases (Figure 4), for all of T cell network outputs. However, the *N* value that gives the highest vulnerability varies depending on the output. For the network outputs ap1, bcat and p70s, we see the maximum vulnerability increase going from single faults to double faults, *N* = 1 and 2, respectively, and reaches a plateau afterwards. For some other network outputs such as cre, nfat, p38, shp2 and sre, the highest maximum vulnerability occurs when *N* = 1. This implies that the functions of these outputs are more dependent to the function of specific individual faulty molecules which are sufficient to induce the extreme network failures at these outputs.

We also prove that the computational complexity, i.e., the running time of the proposed extreme signaling failure analysis algorithm (Methods, Section B) is *O* (*K*^3^), where *K* is the number of intermediate molecules in the network. This efficient algorithm is in contrast with an exhaustive search having an exponential running time, *O* (*K*^*K*/2^), that quickly becomes impractical to implement, as *K* increases. For example, for a network with *K* = 50 molecules, the proposed algorithm complexity is in the order of 50^3^ ≈ 1.3 ×10^5^, which is much smaller than 50^25^ ≈ 3 ×10^42^, the exhaustive search complexity.

The proposed extreme signaling failure algorithm makes use of another proposed algorithm (Methods, Section C) that properly incorporates the effects of signaling feedbacks in the extreme signaling failure analysis. Essentially it determines the number of time points (clock cycles) needed for network analysis and simulation while computing the vulnerability levels to find the extreme signaling failures, so that we prevent performing network simulations longer than what is needed. Usefulness of this algorithm is demonstrated by computing the requited number of clock cycles for vulnerability analysis of the ERBB and the T cell signaling networks (Methods, Sections D and E, respectively).

Overall, the proposed algorithms have the potential to uncover certain aspects of abnormal signaling network behaviors that can contribute to the development of the pathology, and may suggest some new therapeutic strategies in the area of targeted therapy in pharmaceutical industry. This study is particularly important in the context of complex trait disorders with poorly understood molecular sources when more than one molecule is often reported to be involved in the pathogenesis of the disease.

**Key Points**

- The main goal of this research is to quantify the vulnerability of a signaling network to the dysfunction of one or multiple molecules, when the dysfunction is defined as an incorrect response to the input signals.
- An efficient algorithm to identify the extreme signaling failures, defined as a pathological phenomenon that results in the highest probability of network failure, is developed and applied to the two experimentally verified ERBB and T cell signaling networks.
- The results reveal that as the number of concurrently dysfunctional molecules increases, the maximum vulnerability values quickly reach to a plateau following an initial increase, which suggests the specificity of vulnerable molecule (s) involved. A specific number of faulty molecules cause the most detrimental damage to the function of the network. Increasing a random number of simultaneously faulty molecules does not further deteriorate the function of the network.
- Such specific group of molecules whose dysfunction causes the extreme signaling failures of the intracellular molecular networks can elucidate the molecular mechanisms underlying the pathogenesis of complex trait disorders, and can offer new insights for the development of novel therapeutics.

